# scKEPLM: Knowledge enhanced large-scale pre-trained language model for single-cell transcriptomics

**DOI:** 10.1101/2024.07.09.602633

**Authors:** Yang Li, Guanyu Qiao, Guohua Wang

## Abstract

The success of large-scale pre-trained language models in the Natural Language Processing (NLP) domain has encouraged their adoption in genomics and single-cell biology. Developing pre-trained models using the rapidly growing single-cell transcriptomic data helps to unravel the intricate language of cells. However, current single-cell pre-trained models primarily focus on learning gene and cell representations from extensive gene expression data, failing to fully comprehend the biological significance of the gene expression patterns and cell types they identify, which leads to limited interpretability and transferability. We propose scKEPLM, a knowledge-enhanced single-cell pre-training language model integrates a biology knowledge graph into the single-cell transcriptome pre-training process. scKEPLM covers over 41 million single-cell RNA sequences and 8.9 million gene relations. Through parallel pre-training of single-cell transcriptome sequences and genetic knowledge, combined with a Gaussian cross-attention mechanism, scKEPLM precisely aligns cell semantics with genetic information, to learn more accurate and comprehensive representations of single-cell transcriptomes. The introduction of knowledge enhancement has improved the identification of important genes in cells by scKEPLM, and greatly enriched the understanding of cell function and disease mechanism. The scKEPLM model has achieved state-of-the-art performance in more than 12 downstream tasks, including gene annotation, cell annotation, and drug response prediction, demonstrating strong generalization and transferability. Further exploration of the model’s interpretability demonstrates its adaptability to variations in gene expression patterns within cells under various physiological or pathological conditions.

---

Single-cell sequencing technology enables the acquisition of cellular information at the level of individual cells. This approach addresses a critical limitation of traditional sequencing technologies, which analyze groups of cells and, as a result, obscure inter-cellular heterogeneity. By focusing on individual cells, researchers can precisely explore cell heterogeneity and life activities, marking a significant shift from group-based to individual cell analysis. Singlecell RNA sequencing (scRNA-seq) is currently the predominant technique in this field. It encompasses the reverse transcription, amplification, and high-throughput sequencing of an individual cell’s mRNA. This process enables the detailed examination and characterization of single-cell molecular profiles, or cellular expression, across diverse species, tissues, organs, and organisms. The insights gained from single-cell transcriptomic data enable a more refined analysis of disease causes and assist in the precise matching of treatment plans. It is highly valuable for precision medicine, paving the way for developing targeted therapies and personalized treatment strategies.

Recently, large pre-training language models, through extensive pre-training followed by fine-tuning, have significantly advanced the fields of Natural Language Processing (NLP) [1] and Computer Vision (CV) [2]. These models’ ability to learn universal representations of language or visual data has been leveraged in bioinformatics, notably for analyzing large-scale, unlabeled single-cell transcriptome datasets. The structural parallels between natural language and genetic sequences, as exemplified by models like scBERT and Geneformer [3], underscore the potential for such models to enhance our understanding of cellular functions, gene expression patterns, and disease mechanisms. However, transitioning from human languages to the genetic lexicon presents challenges. Despite their complexity, pre-trained models with billions of parameters often struggle to generalize across the diverse tasks of single-cell transcriptomics. Their reliance on unsupervised learning from generic biological sequences can result in models that excel at capturing basic sequence patterns but fall short in interpretability and adaptation to specific biological problems.

A possible solution to these challenges is to incorporate fundamental biological knowledge into both the pretraining and fine-tuning stages. By embedding pre-training models with established genetic principles, researchers can steer them toward a more sophisticated grasp of genetic semantics and cellular dynamics. Such an enriched foundation not only improves the models’ interpretability but also improves their ability to transfer important knowledge for downstream tasks. Integrating genetic knowledge upfront is thus pivotal for unleashing the full potential of pre-training language models in navigating the intricate landscape of single-cell transcriptomics.

In this work, we propose a knowledge-enhanced large-scale pre-training language model scKEPLM (Fig.1) based on 41 million single-cell RNA sequencing data. The scKEPLM model integrates a genetic knowledge graph comprising 8.9 million gene relations into the pre-training stage for the first time and simultaneously pre-trains single-cell transcriptome sequences and genetic knowledge.

**Fig. 1:**
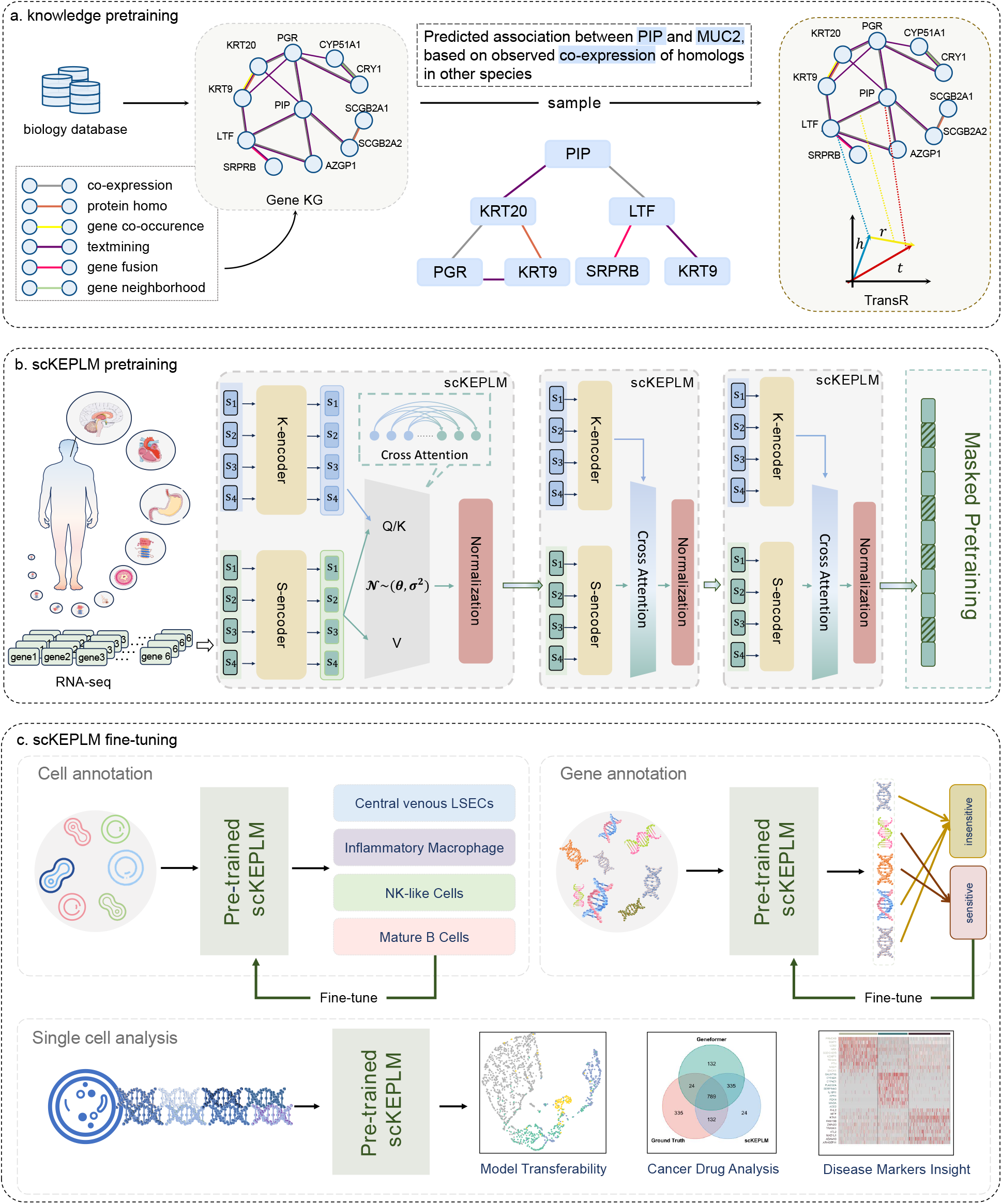
An illustrative diagram of scKEPLM. **a**, Genetic knowledge graph pre-training. The TransR method is employed to learn valuable initial embeddings for genes and their interactions from the genetic knowledge base STRING. It captures the inherent structure and semantics of the knowledge graph, offering knowledge enhanced representations for genes within the biological context. **b**, Pre-training phase of scKEPLM. The pre-training stage of scKEPLM is based on a Masked Language Model (MLM) architecture, which is commonly used in natural language processing tasks. In the context of scRNA-seq, gene expression sequences are regarded as sentences, with each gene or transcript corresponding to a word in the biological language, and the expression level of the gene representing the “meaning” of the sentence. scKEPLM leverages MLMs to predict missing or masked elements in these sequences, enabling it to understand the contextual relationships between genes and infer the patterns of gene deletions. scKEPLM consists of two parallel encoders: the sequence encoder and the knowledge encoder. These encoders are combined and standardized using a Gaussian cross-attention mechanism, ultimately encoding rich genetic knowledge in the RNA sequences. **c**, Fine-tuning phase of scKEPLM. It enables the model to leverage the generalized semantic knowledge acquired during pretraining, enhancing its ability to comprehend and perform well on specific downstream tasks, particularly when there is limited labeled data available. During the finetuning stage, we designed 12 downstream tasks, including both cell and gene levels, to demonstrate scKEPLM’s generalization ability across various tasks.

scKEPLM marks significant advancements and discoveries in four aspects of single-cell gene expression analysis. Firstly, it introduces a novel knowledge-enhanced framework that boosts unsupervised single-cell RNA sequence pre-training. The integration of biological priors leads to significant improvements in crucial bioinformatics tasks, such as gene and cell annotation. scKEPLM incorporates genetic principles with unlabeled transcriptome data during its training phase, alongside a comprehensive pre-training corpus. It emphasizes the critical importance of data diversity and volume in improving the model’s generalizability. Secondly, the scKEPLM demonstrates exceptional generalization and domain adaptation capabilities. In the task of cell annotation, through pre-training on sequencing data from various tissues and platforms, followed by cross-domain fine-tuning, the scKEPLM demonstrates its robust adaptability and flexibility. The cross-platform compatibility opens new avenues for integrated analysis across various platforms, potentially accelerating the integration of multi-source data and broadening the application of single-cell technologies in a wider range of fields. Thirdly, the scKEPLM demonstrates heightened sensitivity in identifying disease-associated biomarker genes when applied to single-cell samples derived from diverse pathological conditions. It highlights the pivotal role of pre-training methodologies in advancing the frontiers of disease research. Finally, scKEPLM employs a Gaussian attention mechanism to model the complex high-dimensional interaction between single-cell semantic expression and biological knowledge. It allows for the precise localization of differentially expressed genes in various cell states. The Gaussian attention mechanism outperforms traditional methods by capturing subtle biological signals embedded in gene expression profiles. This advancement in detecting biological functions and gene expression variations related to pathological conditions significantly enhances our ability to unravel complex biological regulatory networks. The efficiency and accuracy of scKEPLM deepen our understanding of the molecular dynamics involved in disease processes.

## 1 Results

### 1.1 Overview of scKEPLM

Single-cell RNA sequencing (scRNA-seq) technologies have revolutionized biology and medicine by enabling the characterization of the transcriptomes of individual cells, thus providing a high-resolution view of cellular diversity and function within tissues and organs. The technological leap has led to the identification of novel cell types, understanding of cellular differentiation processes, and insights into the mechanisms of disease at an unprecedented level of detail. Single-cell RNA transcriptomics pre-trained models aims understanding complex biological systems at the single-cell level.

Our proposed model scKEPLM, as shown in Fig. 1, consists of three main components: Genetic knowledge pre-training, scKEPLM pre-training and fine-tuning in downstream tasks. scKEPLM takes a significant volume of unsupervised single-cell RNA sequences as input. Incorporating biological knowledge is essential for enhancing the performance of large pre-trained models [1, 4, 5]. To accomplish this, we initially apply an unsupervised graph pretraining method on a genetic knowledge graph to capture a foundational gene knowledge representation. We select the STRING database (Search Tool for the Retrieval of Interacting Genes/Proteins) as an external knowledge graph as it provides critical protein-protein interaction data, supports pathway analysis, and aids in identifying disease biomarkers for diagnosis and treatment [6, 7]. During the pre-training phase, scKEPLM employs multiple pre-training units. Each encoding block consists of an RNA sequence encoder module (S-encoder), a knowledge graph encoder module (K-encoder), and a Gaussian cross-attention layer. Similar to the BERT model, scKEPLM uses a masking strategy for pre-training. It masks 15% of the input single-cell transcriptomic data and uses the unmasked data to predict the hidden positions. After unsupervised pre-training on a large volume of unlabeled single-cell transcriptomic sequencing data, scKEPLM is fine-tuned for various gene and cell-level tasks, enabling the successful application of the pre-trained model to complex downstream tasks.

We construct a comprehensive dataset for the pre-training of scKEPLM, comprising 41.83 million single-cell transcriptomes from 30 major tissues, encompassing over 60 cell types, derived from publicly available datasets. To ensure the quality of the data, scalable filtering criteria were applied to exclude cells with high mutation burdens or evidence of damage, effectively minimizing the influence of noise [3].

### 1.2 scKEPLM boosts the performance of gene annotation

We designed a series of experiments to evaluate the performance of the pre-trained scKEPLM model on three primary gene annotation tasks. To fine-tune the model, we selected a limited dataset for each task, and evaluated the model’s robustness at the gene level. We compared the scKEPLM model with a variety of typical machine learning methods as well as several state-of-the-art pre-trained models. To avoid overfitting, all experiments were subjected to five-fold cross-validation, and the average results across the folds served as the final performance metric.

#### 1.2.1 Gene dosage sensitivity identification

Previous studies have indicated that the dosage sensitivity of specific genes plays a critical role in the pathogenicity associated with copy number variations (CNVs) in humans[8]. It is also a crucial determinant in evaluating the penetrance of gene mutations and understanding the mechanisms underlying genetic diseases. Therefore, determining the dosage sensitivity of genes is essential for identifying the pathogenic factors in human diseases [8, 9]. Despite existing methods such as eQTL and its variants, which use allele frequencies to identify gene dosage sensitivity, they have not accurately captured transcriptional dynamics. This is due to the constant effect size values, which do not account for changes in cellular states. Furthermore, gene dosage sensitivity is influenced by gene function, meaning that changes in gene dosage can affect different tissues in distinct ways.

Using previously validated dose-sensitive and dose-insensitive genomic datasets as the gold standard, we finetuned the pre-trained scKEPLM model using 10,000 randomly selected single-cell transcriptome data samples. We benchmarked the performance of scKEPLM against other methods for identifying gene dosage sensitivity, based on the fine-tuned scKEPLM. The scKEPLM model demonstrated superior prediction accuracy, with an impressive area under the curve (AUC) score of 0.936-2% higher than the SOTA model Geneformer (Fig. 2a).

**Fig. 2:**
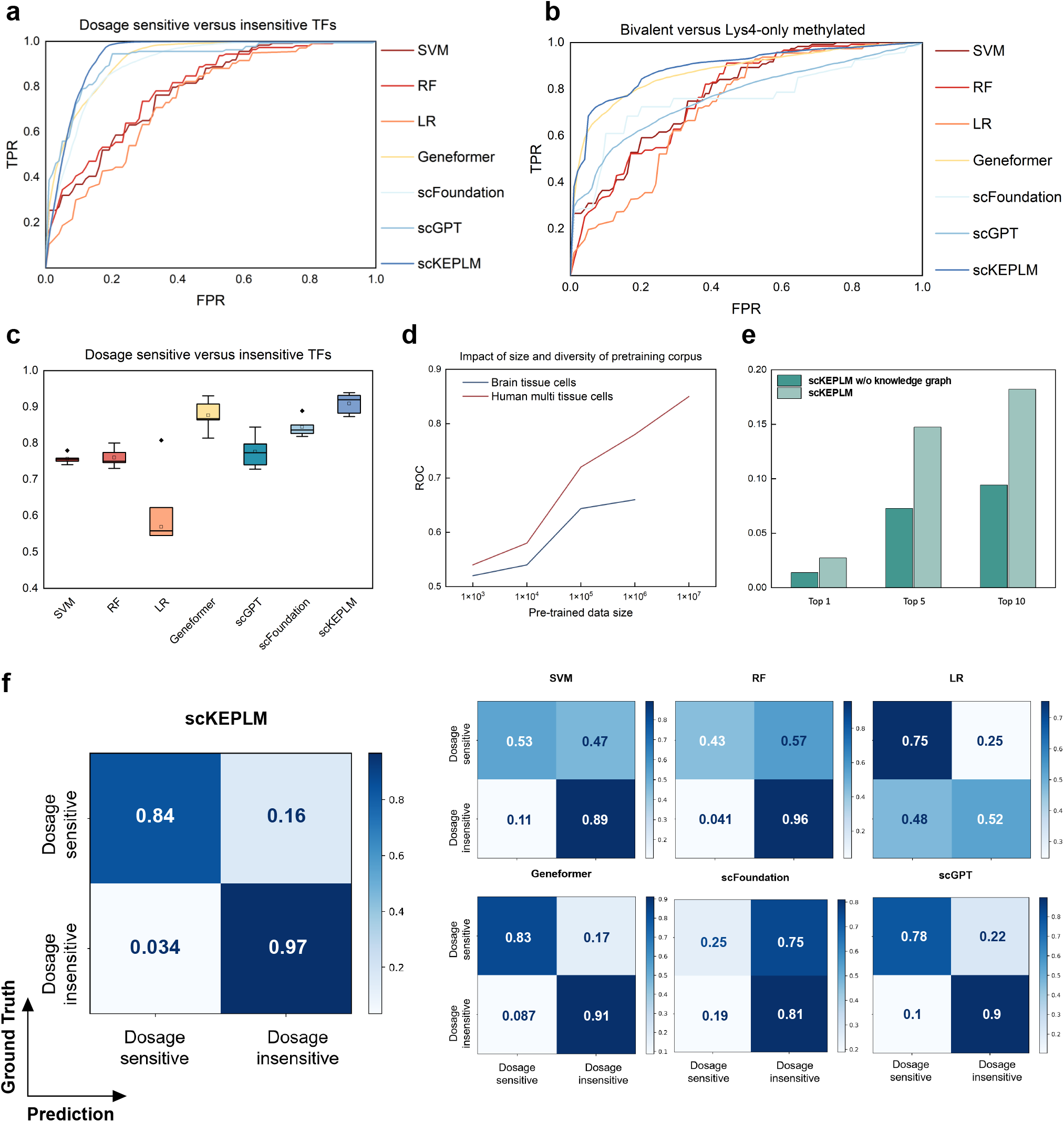
scKEPLM for efficient gene annotation. **a**, Gene dosage sensitivity ROC curve performance comparison. The models are fine-tuned with limited data to differentiate the dosage sensitivity of transcription factors. Compared with the traditional or current state-of-the-art methods, scKEPLM significantly outperformed other approaches. **b**, Comparison of ROC curves for bivalent and Lys4-only Methylation prediction. **c**, Comparison of gene dose sensitivity classification F1-scores using box plots. **d**, Evaluation of the impact of pre-training data scale on the generalization performance of the scKEPLM model. We selected the “brain tissue cells” and “human multi-tissue cells” from scKEPLM-41M to examine the impact on the model’s generalization capability. **e**, To compare knowledge-enhanced (scKEPLM) and non-knowledge-enhanced (scKEPLM w/o knowledge graph) pretrained models, the bar chart shows the ratio of similar marker genes within the top k. The x-axis represents the Top k (the top 1, 5, 10 most similar genes), and the y-axis represents their proportion in the total dataset. **f**, Confusion matrices of the six comparision methods and scKEPLM model.

The box plot evaluating F1-score for classifying dosage-sensitive and insensitive transcription factors (Fig. 2c) highlighted the scKEPLM model as the top performer. It achieved the highest median F1-score with a narrow range, highlighting its consistent superiority due to knowledge encoding. In contrast, the Geneformer model, despite a high F1-score, exhibited greater variability, suggesting potential inconsistency in different scenarios. Traditional machine learning methods like SVM (Support Vector Machine), RF (Random Forest), and LR (Logistic Regression) underperformed compared to Transformer-based models. SVM and RF were more reliable than LR. Overall, the data conclusively showed that the knowledge-enhanced scKEPLM model outperformed all others in this task.

The visualization analysis of the confusion matrix further demonstrated the above results (Fig. 2f). It showed that scKEPLM had a higher recall rate and precision in identifying dosage-sensitive samples, while also exhibiting extremely high specificity in dosage-insensitive samples. Although the overall accuracy of the Geneformer model was close to that of scKEPLM, the latter outperformed in terms of specificity, false-positive rate, and false-negative rate. The scFoundation [10] model might have introduced too much gene-level noise in the process of perturbation of its model depth perception indicators, leading to a higher false-positive rate in its predictions. These findings not only demonstrate the advantages of scKEPLM in accuracy but also highlight the importance of its predictive capability and biological insights when used to predict dose sensitivity.

#### 1.2.2 Identification of bivalent chromatin structure

The bivalent chromatin structure present in embryonic stem cells is crucial for cell differentiation [11]. This distinctive chromatin structure harmonizes gene silencing and activation in undifferentiated cells, preserving a delicate balance for genes pivotal to cell differentiation and developmental processes [12]. The bivalent chromatin structure in embryonic stem cells is associated with both activating (H3K4me3) and repressive (H3K27me3) epigenetic marks, and their interaction determines the fate of embryonic stem cells and affects whether they differentiate or remain undifferentiated.

We randomly selected 15,000 embryonic stem cells [13] from the panglaoDB database to fine-tune the pretrained scKEPLM model, to identify genes marked with bivalent epigenetic signals from those that only have unmethylated or H3K4me3-related promoter regions. The ROC curves showed the performance of different models in differentiating bivalent genes from those exhibiting Lys4-only methylation. The closer a curve was to the top left corner, the better the model’s performance (Fig. 2b and Extended Data Fig. 1a-b). Geneformer and scKEPLM demonstrated the best performance among the evaluated models, as their ROC curves were closest to the top-left corner, indicating a higher true positive rate (TPR) for a given false positive rate (FPR). The SVM and RF models exhibited comparable performance, with SVM slightly outperforming RF overall. The LR model demonstrated the lowest performance, with its ROC curve positioned the farthest from the top-left corner, indicating a reduced true positive rate (TPR) at equivalent false positive rate (FPR) levels. The significant performance of scKEPLM can be attributed in part to its ability to fine-grained information about gene-gene interactions through the gene knowledge graph, thereby enhancing the model’s understanding of biological processes. As a result, it is able to outperform other SOTA models, like Geneformer, in gene annotation tasks. In contrast, traditional machine learning models often struggle to assimilate contextual knowledge, resulting in poor generalization and weaker performance in gene annotation.

Overall, the strengths of the scKEPLM model can be attributed to its design, which is highly integrated with biological signatures, making it more persuasive in predicting stem cell behavior and understanding the epigenetic mechanisms involved in embryonic development. The successful application of this model underscores the importance of combining deep learning with biological intuition in the analysis of biological data.

#### 1.2.3 Identification of long and short-range transcription factors

Genomic distance is crucial for explaining regulatory variation and inferring target genes from transcription factor genomic occupancy data. Previously, other researchers systematically integrated thousands of spectra of transcription factor binding and histone modifications using Chromatin Immunoprecipitation Sequencing (ChIPseq), and analyzed gene expression profiles of thousands of genes to identify two types of transcription factors with differing regulatory ranges [14].

We fine-tuned scKEPLM using single-cell transcriptomes from approximately 12,000 cells undergoing iPSC [15] to cardiomyocyte differentiation to differentiate the long-range and short-range transcription factors, without using associated ChIP-seq or genomic distance data (Extended Data Fig.1b-d). Compared to other methods, scKEPLM significantly enhanced the predictive capability for the regulatory scope of transcription factors, whereas results from other methods were almost random. Therefore, the fine-tuned scKEPLM model not only captures transcription factor features within regulatory ranges but also excels in identifying complex features from transcriptomic data alone.

The results highlight the model’s strengths in identifying and differentiating between long-range and shortrange regulatory factors. The scKEPLM model excels by learning intrinsic patterns within transcriptome data and understanding the contextual relationships of genes, thereby overcoming the limitations of traditional datasets and delineating the extent of regulators’ influence. These traits suggest that scKEPLM possesses significant potential for predicting cellular metabolic regulation and gene regulatory networks, offering fresh insights into disease mechanisms and potential targets for gene therapy. In contrast, Geneformer, despite being a state-of-the-art pretrained model for single-cell transcriptome data, has limitations in modeling the deep semantic relationships between genes and is highly sensitive to data noise, which hinders its ability to fully capture the complexity of regulatory networks.

#### 1.2.4 Effects of quantity and diversity of pre-training data

The quantity and diversity of pre-training data significantly influence the performance and generalization capabilities of the pre-training model. We tested the robust performance of the scKEPLM model concerning pre-trained data by evaluating its sensitivity to gene dosage in experiments using datasets of varying sizes. In the experiments, we focused on two different types of cells: “Brain Tissue Cells” [16] and “Human Multi-Tissue Cells”. By varying the scale of the pre-training data from 1,000 to 10,000,000 on a logarithmic scale and observing the corresponding ROC values (*y*-axis from 0.5 to 1.0), we discovered that the performance of both cell types in the gene dosage sensitivity identification task improved as the scale of the pre-training data increased (Fig. 2d).

However, compared to “Brain Tissue Cells”,”Human Multi-Tissue Cells” started at a higher performance level, and their performance increased more noticeably as the data size expanded. When the pre-training data contains a more diverse range of tissue types, scKEPLM is able to learn and generalize knowledge more effectively (Fig. 2d). This indicates that for handling multi-tissue cell data, scKEPLM might necessitate larger datasets to achieve peak performance. With 1 million samples from the pre-training data, we noted that the ROC performance score for “Human Multi-Tissue Cells” approached 0.784, whereas the score for “Brain Tissue Cells” was approximately 0.661, significantly lower than the former.

In addition, we conducted ablation experiments on the scKEPLM model to verify whether the inclusion of the knowledge graph enhances the model’s ability to identify intracellular gene associations. We compared the scKEPLM model with its degenerate version, scKEPLM w/o knowledge graph, in the acquisition of single-cell representations and the identification of marker genes in single cells. The cosine similarity score is a metric to measure the correlation between two genes, with higher scores indicating a closer spatial relationship in their embeddings. Without fine-tuning, using the Muraro dataset and particularly focusing on *α*-cells as our test dataset, we evaluated the results of the two models in evaluating the cosine similarity of gene embeddings within cells. As shown in Fig. 2e, the performance of the model with a knowledge graph in detecting the 1, 5, and 10 most relevant genes was nearly twice that of the model without a graph, indicating that the addition of a knowledge graph significantly enhanced the model’s ability to recognize subtle associations between genes. Hence, the knowledge graph, by providing a rich background knowledge framework, enriched the model’s comprehension and excavation capabilities regarding complex biological signals, thus offering new directions for the exploration of disease mechanisms and the development of biotechnology. Meanwhile, this high degree of relevance in gene embeddings might enhance the model’s sensitivity to differences between cells, leading to exceptional performance in cell annotation tasks.

### 1.3 scKEPLM demonstrates outstanding performance in cell annotation tasks

In biology, cells exhibit distinct phenotypes, diverse cell surface antigens, and unique gene expression profiles [17]. The variations enable cells to perform varied roles in human genetics and development. Single-cell annotation, which involves analyzing transcriptomic data from individual cells, assists in identifying and categorizing different cell types [18, 19]. The process uncovers the cellular composition of tissues and organs, improving our understanding of cell functions, differentiation stages, metabolic activities, and signaling networks. Therefore, we fine-tuned the scKEPLM model and established a connection between single-cell transcriptome data and cell types and annotations to evaluate scKEPLM’s capacity to represent cells and their intricate functions.

#### 1.3.1 Cell annotation and cell subtype annotation

To demonstrate the capability of the scKEPLM pre-trained model in learning cell-level representations (Fig. 3a), we evaluated its performance in cell annotation and compared it to leading deep learning models as benchmarks across various tissues. The remarkable performance of the scKEPLM pre-trained model in cell annotation tasks not only reflects its strong learning ability across various tissue cell types but also highlights its superiority over currently leading deep-learning models, such as scDeepSort [20] and CaSTLe [17]. Although these models have made some progress, their generalization ability is limited in environments with scarce or poor-quality data, as they rely on a large amount of labeled data. The advantage of scKEPLM lies in its use of a significant amount of unlabeled data for training through self-supervised methods, rather than relying solely on annotated training samples. scKEPLM is particularly outstanding in tests of immune and spleen cells, with accuracy and F1-score almost always leading other models, indicating a deeper understanding of gene expression patterns across different tissue types and an ability to capture subtle differences. Geneformer also showed good F1-score in specific tissue types such as liver, lungs, and pancreatic cells. scDeepSort, however, generally performed poorly, especially in the annotation tasks of kidney and pancreatic cells, with accuracy and F1-score far behind other models, possibly due to inadequate generalization ability or less precise capturing of gene expression patterns in specific cell types.

**Fig. 3:**
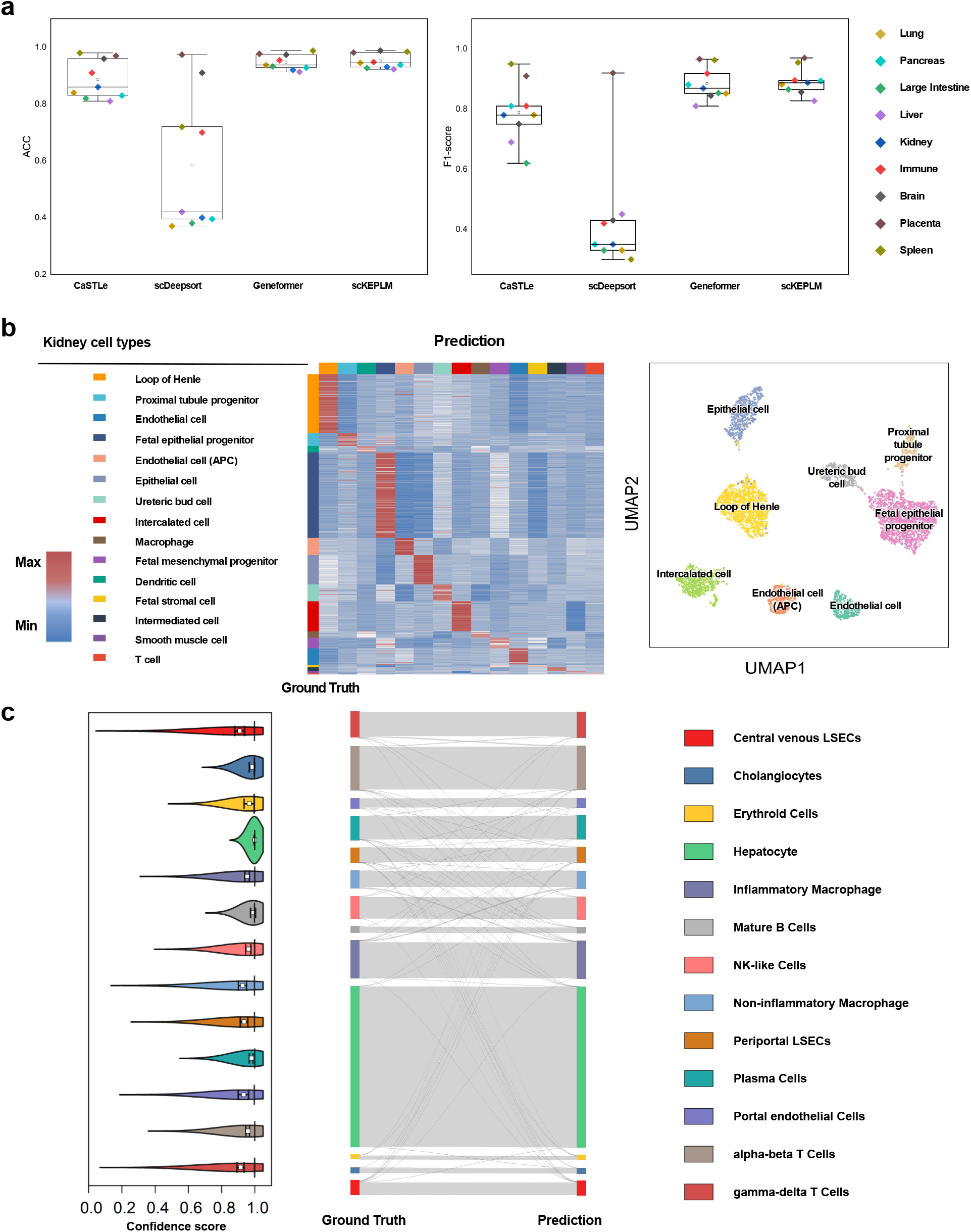
Cell annotation using scKEPLM. **a**, The performance of scKEPLM in cell annotation. The scKEPLM, when fine-tuned with a limited amount of data, outperforms pre-training methods like Geneformer and other deep learning methods in the metrics of Accuracy and F1-score. **b**, Heatmap and UMAP Visualization of scKEPLM on Kidney Cells. In the heatmap, red color signifies higher predictive scores, while blue represents lower scores. The UMAP visualization displays cell type annotations on the kidney dataset. Circular symbols in the legend represent cell types in the UMAP plot, while square symbols correspond to cell types in the heatmap. **c**, Sankey diagram for cell annotation tasks on the Zheng68k dataset.

The heatmaps visually demonstrated the relationship between the actual kidney cell types and the predictions made by the scKEPLM model (Fig. 3b and Extended Data Fig. 2-3). The depth of color served as a measure of match, with darker colors representing higher consistency, meaning that the model’s predictions of cell types highly matched the actual categories; conversely, lighter colors indicated lower prediction accuracy. From the heatmap, when the color tends towards darker on the diagonal, it indicates highly accurate cell type predictions in the corresponding positions. Furthermore, the cell representations learned by the scKEPLM model were visualized using UMAP technology. As shown in Fig. 3b, specific types of cells clustered together in the graph, with clear boundaries visible between different groups of cell types. This distinct clustering and differentiation between clusters proves that the scKEPLM model can not only recognize cell types but also capture the complex relationships and differences between cells. Mapping high-dimensional data to a low-dimensional space, UMAP has demonstrated the high accuracy of scKEPLM in learning cell representations. These findings not only confirm the model’s effectiveness but also offer a robust tool for future cell classification and biomedical research through unsupervised learning.

Furthermore, we conducted a lightweight fine-tuning of scKEPLM to evaluate its performance in cell and cell subtype annotation tasks. In the comparison of over ten competitive approaches on four public datasets, we found that scKEPLM continues to perform exceptionally well, even with fine-tuning restricted to only the last layer. From the sankey diagram in Fig. 3c and Extended Data Fig. 4a-b, we can conclude that various models, including scKEPLM, scBERT [21], scNym [22], SciBet [23], Seurat [24], SingleR [25], CellID (CellID cell and CellID group) [26] and scmap (scmap cell and scmap cluster) [27], displayed notable performance differences across diverse datasets. scKEPLM demonstrated high consistency across all the tested datasets Extended Data Fig. 4), indicating superior generalizability.

#### 1.3.2 scKEPLM exhibits exceptional generalization capabilities

Transcriptomic data from the real world often come from different sequencing platforms and technologies. Variations in sequencing depth, read length, and error rates among different sequencing technologies lead to differences in the sensitivity and accuracy of gene expression detection, as well as in the resulting gene expression profiles. Here, we attempted to evaluate whether the cell representations learned by scKEPLM possess domain adaptability, that is, the stability of cell annotation across sequencing from different platforms, and to verify if the cell annotations remain consistent in different sequencing data from the publicly available datasets of iPSC [15] differentiation into cardiomyocytes on the Drop-seq (single-cell) and DroNc-seq (single-nucleus) platforms. Initially, we fine-tuned the scKEPLM model on the Drop-seq data and performed cell annotation separately on different platforms (Fig. 4a-b). The fine-tuned model exhibited outstanding performance in the cell annotation task. Subsequently, we tested whether fine-tuning the model with data obtained from Drop-seq sequencing would affect its ability to generalize cell type distinction in another set of cells sequenced on the DroNc-seq platform. The cell embeddings from the fine-tuned scKEPLM were not affected by the sequencing technology and continued to cluster by cell type.

**Fig. 4:**
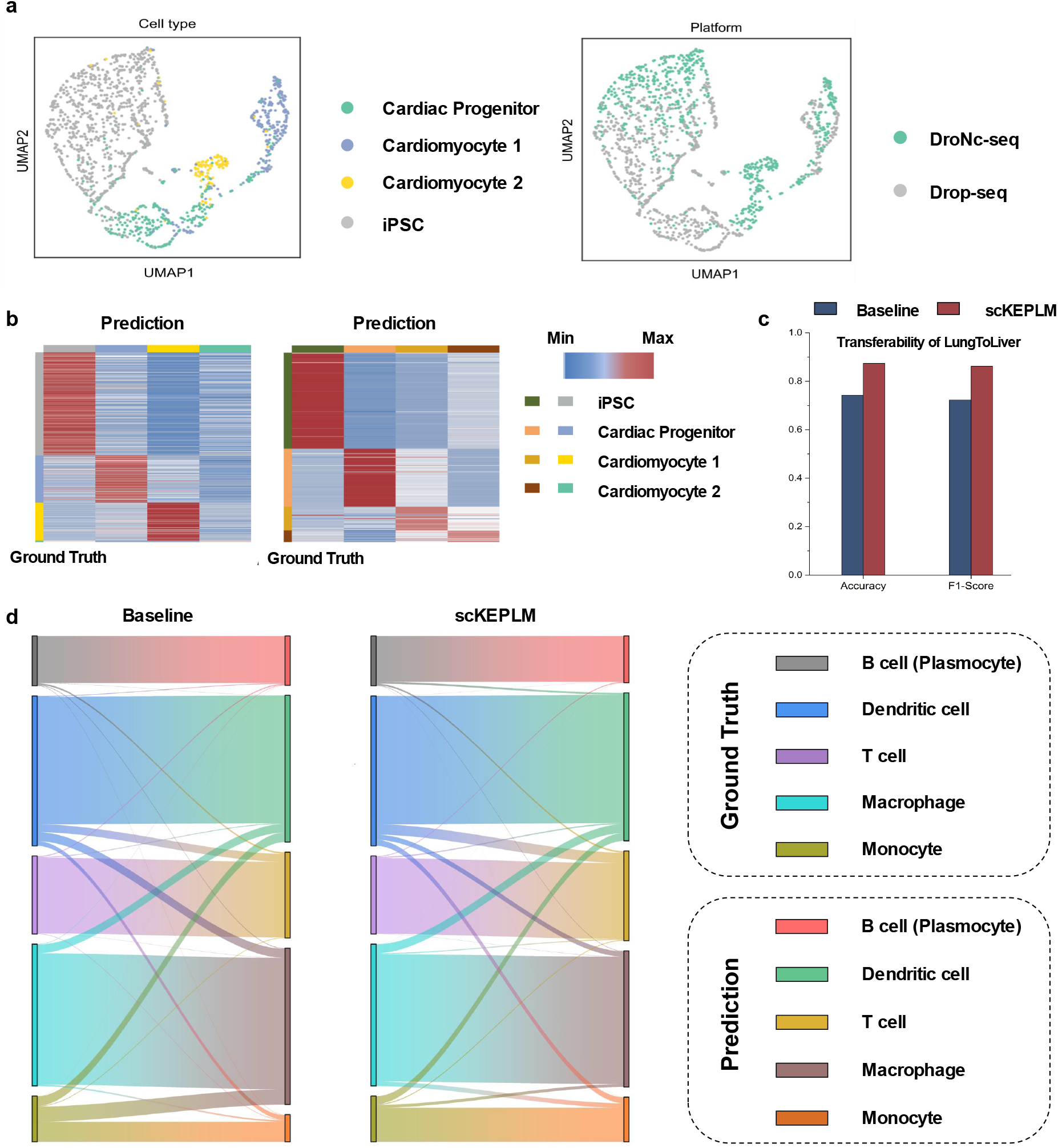
Domain adaptability of scKEPLM. **a**, Fine-tuning scKEPLM with limited single cell sequences obtained from the Drop-seq platform, we can examine the UMAP visualization results for annotated Drop-seq (which did not participate in the fine-tuning) and cells obtained from the DroNc-seq platform. The experimental results demonstrate that the cell representations learned by scKEPLM from different sequencing platforms can be automatically clustered by cell class, regardless of the platform. It highlights the cross-platform representation learning ability of scKEPLM. **b**, The heatmaps illustrate the results of cell annotations by the scKEPLM model that was fine-tuned on Drop-seq data, applied to both Drop-seq and DroNc-seq datasets. **c**, Bar chart of evaluation metrics for scKEPLM across tissues. **d**, Sankey diagram of scKEPLM in cross-tissue cell annotation.

In addition, to test scKEPLM’s cross-tissue cell annotation ability, we fine-tuned the model using liver data and subsequently applied the fine-tuned model to predict annotations for five distinct cell types from lung tissue. As shown in Fig. 4c, the scKEPLM model outperformed the Geneformer model in terms of precision and F1-score. The impressive cross-tissue generalization performance of the scKEPLM model might be attributed to its learning mechanism that integrates prior knowledge, providing the model with a more accurate and extensive capability for cell characteristic recognition compared to conventional pre-training methodologies. Guided by a knowledge graph, scKEPLM can classify cell types more accurately, especially when dealing with complex tissue features and cellular functional changes. This demonstrates its powerful generalization and adaptability with this enhanced pre-training strategy.

Additionally, to further analyze the results, we employed a Sankey diagram to compare the predictions of the scKEPLM model with those of the baseline model (Fig. 4d). We found that the macrophage cells exhibited poorer performance in the baseline model. Although Geneformer can learn from a large amount of data, they lack the direct utilization of biological knowledge. This might result in the models’ inability to accurately capture the subtle differences when cell types exhibit significant heterogeneity across different tissues. For instance, in predicting macrophage cells, ordinary pre-training models might not have sufficient capability to identify and distinguish the specific changes in macrophages under different biological environments.

### 1.4 scKEPLM enhances cancer drug response prediction

Analyzing drug screening data from diverse gene expression databases enables researchers to identify optimal clinical strategies for oncology drugs. The introduction of single-cell RNA sequencing (scRNA-seq) has significantly advanced this field, allowing for the exploration of the heterogeneity in cancer cell subgroup responses to drugs, thereby potentially enhancing treatment efficacy.

We integrated scRNA-seq drug response datasets for Erlotinib, Cisplatin, and Docetaxel, as previously reported [28]. These datasets provided detailed drug response annotations for individual cells, indicating their sensitivity or resistance. Utilizing these datasets, we fine-tuned and compared the scKEPLM model against the Geneformer model for predicting drug sensitivity and resistance. As shown in Fig. 5a, the scKEPLM model excelled in predicting responses to all three drugs, outperforming the SOTA model. It demonstrated scKEPLM’s ability to capture and interpret biological signals to understand complex drug responses. In the Cisplatin test, the scKEPLM achieved a nearly 8% enhancement in AUC performance, emphasizing its accuracy.

**Fig. 5:**
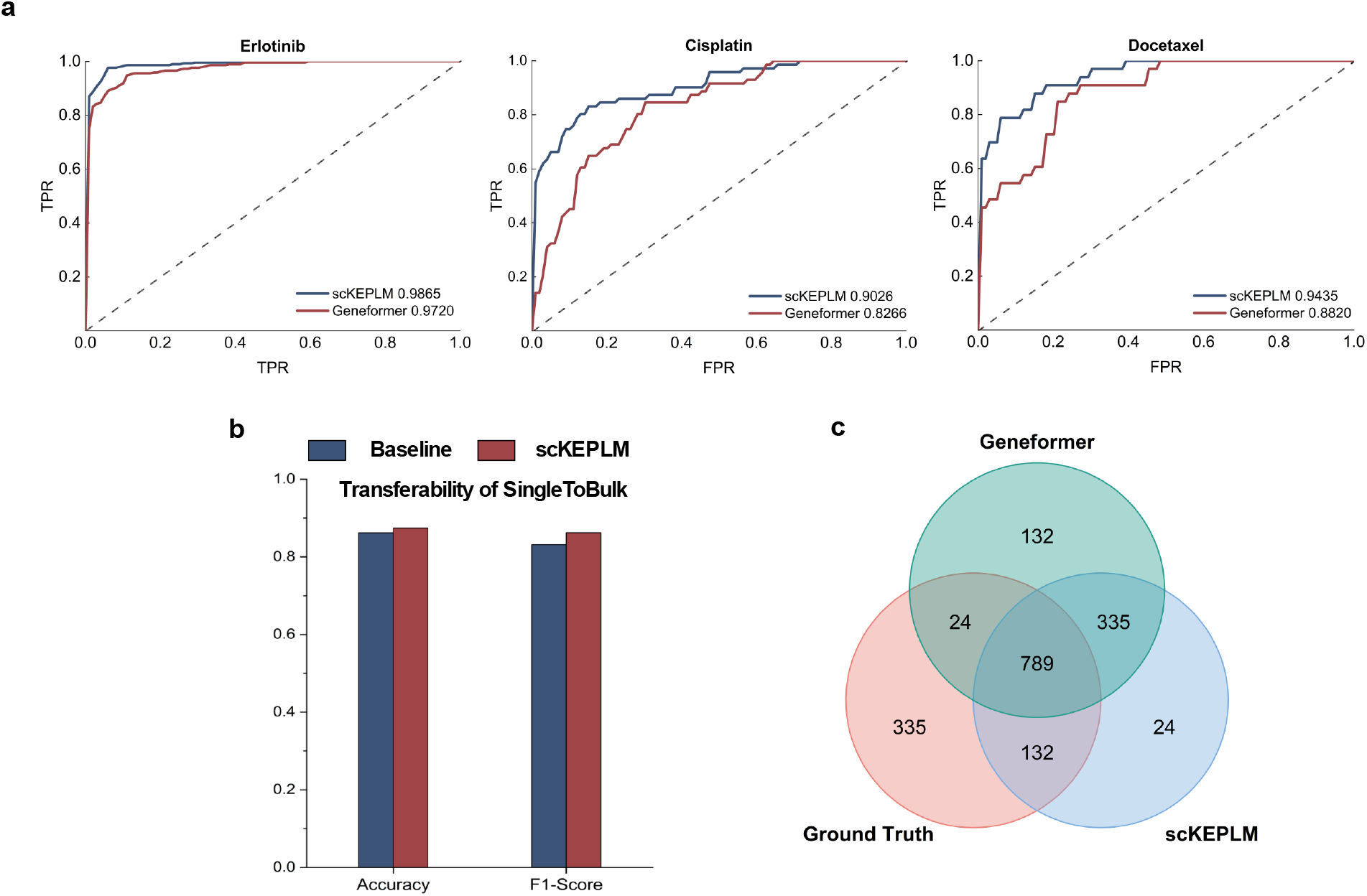
Cancer drug response prediction by scKEPLM. **a**, ROC curves for the predictive performance of scKEPLM and baseline models on three drugs: Erlotinib, Cisplatin, and Docetaxel. **b**, Fine-tuning results of scKEPLM on single-cell data including accuracy and F1-score on bulk level data. **c**, Venn diagram of the prediction results by scKEPLM and baseline models, fine-tuned on single-cell data and on bulk RNA-seq data.

We fine-tuned the scKEPLM model using scRNA-seq data specific to Docetaxel and then applied it to bulk RNA-seq data for predictions, to further validate the model’s generalization capability. As illustrated in Fig. 5b, under the evaluation metrics of accuracy (ACC) and F1-score, compared to the baseline model, the fine-tuned scKEPLM showed a 3% increase in ACC and demonstrated a higher F1-score, indicating the model’s effectiveness in predicting drug responses cross different data levels. Additionally, we constructed a Venn diagram (Fig. 5c) that highlights the 108 additional samples correctly predicted by scKEPLM compared to the baseline model. These results validate the model’s ability to analyze single-cell data and identify genes associated with drug responses.

The aforementioned results emphasize the scKEPLM model’s broad applicability and accuracy in the realms of drug screening and cancer research. By incorporating genetic insights, scKEPLM offers a robust tool for celllevel analysis, enabling a deeper comprehension of drug interactions with various tumor cells and their impact on cell survival and proliferation. This capability is instrumental in guiding the development of future personalized treatment strategies.

### 1.5 scKEPLM exhibits strong interpretability

To investigate how the scKEPLM model captures cell features after fine-tuning, and whether it can identify marker genes in abnormal cells of diseased patients, we conducted a Gaussian weight analysis on the scKEPLM model. During fine-tuning, the optimization of marker data led to the Gaussian cross-attention assigning higher weights to certain genes. These weights can reflect the uniqueness of certain genes within cells and which genes are more important, thereby providing a biological interpretation for the model.

Our proposed model, scKEPLM, has a three-layer architecture, each layer equipped with a weight allocation strategy that merges sequence and knowledge encoding through a Gaussian cross-attention mechanism. This strategy differentially allocates weights, assigning higher values to important genes and lower values to less significant ones, thereby accentuating the distinctions among them. The primary goal of this approach is to improve the model’s semantic analysis capabilities and environmental adaptability. It enables the model to more efficiently learn representations of cells and genes in unlabeled scenarios.

#### 1.5.1 Insights into disease marker genes by scKEPLM

Heart failure is increasingly receiving attention in the realm of public health, manifesting itself through a collection of clinical features, chiefly characterized by impaired cardiac contractile function. Previous studies have mainly focused on changes in transcription and protein expression in failing hearts, yet often overlooked the molecular changes of uncommon cell types in analyses. We collected nearly 560,000 single-cell RNA sequencing (scRNA-seq) data [29] from the left ventricles of 11 dilated cardiomyopathy (DCM) hearts, 15 hypertrophic cardiomyopathy (HCM) hearts, and 16 non-failing (NF) hearts to test whether the scKEPLM model could differentiate gene anomalies in normal human cells from those in disease cells.

By fine-tuning scKEPLM, we aimed to differentiate non-failing (normal) myocardial cells from DCM and HCM disease cells. On this basis, we constructed a list of marker genes for DCM and HCM disease cells and used Gaussian weights from the scKEPLM model’s prediction process as an indicator to evaluate the importance of these genes. Notably, these marker genes generally received higher weights in the fine-tuned scKEPLM model (as shown in Fig. 6a-b and Supplementary Table S1), demonstrating the model’s accuracy in capturing gene expression patterns in diseased cells. For example, mutations in the TNN gene can lead to structural or functional abnormalities in myosin-binding protein C [30], affecting the normal contractile function of the myocardium. During this process, the weights generated by the fine-tuned scKEPLM model were significantly higher than those in normal cells, highlighting the model’s ability to learn specific disease biomarkers, their interactions, and roles in biological pathways, thereby offering a more comprehensive understanding of disease mechanisms.

**Fig. 6:**
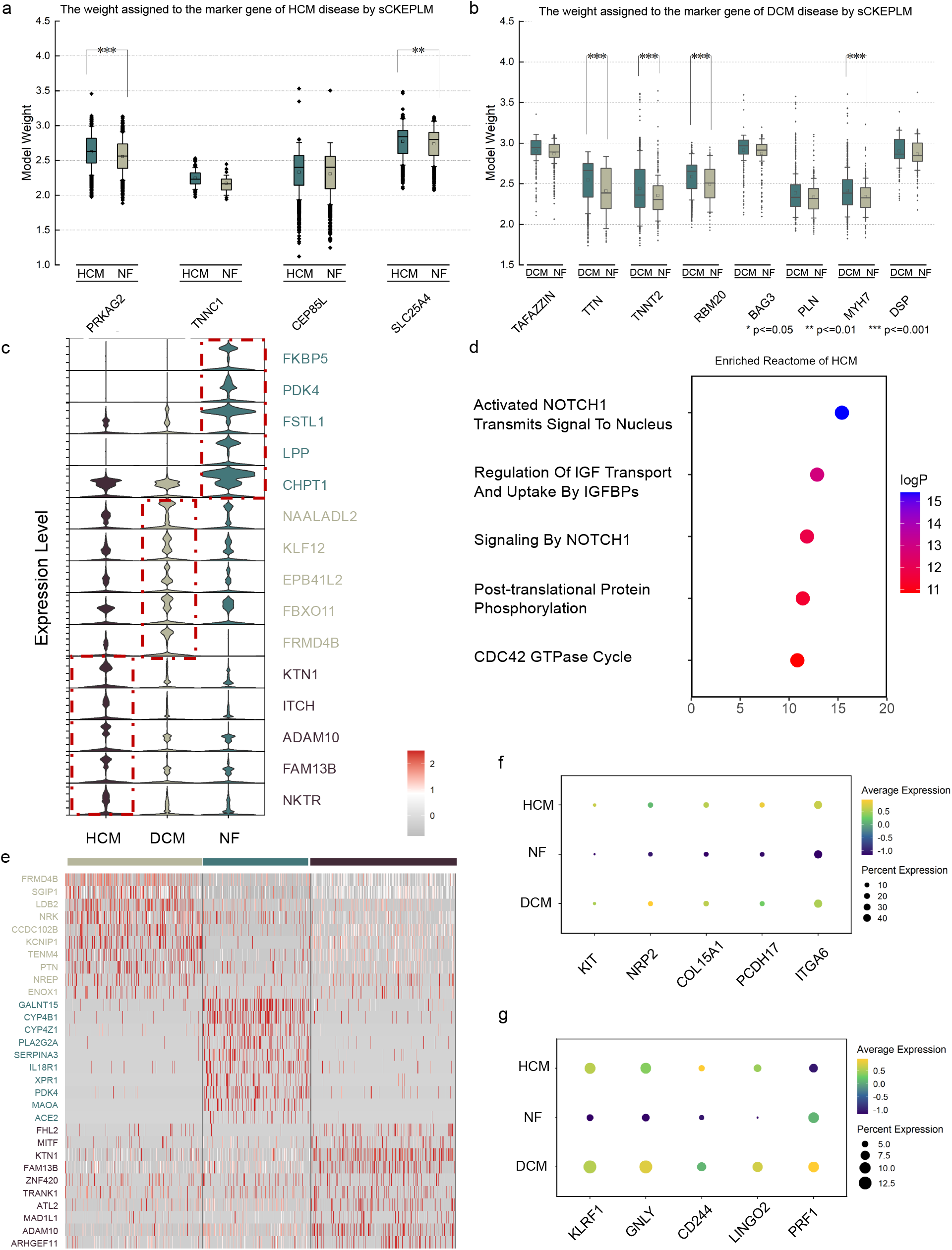
Insights into disease marker genes by scKEPLM. **a**, Boxplot of marker gene weight distribution for HCM patients and a normal population for HCM genes. **b**, Boxplot of marker gene weight distribution for DCM patients and a normal population for DCM genes. **c**, Violin plot for differential analysis of gene weights in HCM, DCM, and normal individuals, selecting the top 5 genes for each group. **d**, Results of the reactome enrichment analysis for the top 5 genes of HCM. **e**, Differential gene heatmap of fibroblasts. **f**, The average expression weight of endothelial cell markers, including KIT, NRP2, COL15A1, PCDH17, and ITGA612 across the NF, HCM and DCM. **g**, The average expression weight of lymphocyte markers, including KLRF1, GNLY, CD244, and PRF1, across the NF, HCM and DCM.

Furthermore, we performed differential analyses on the weights the genes confer on cells in samples from the three populations. Fig. 6c and Supplementary Table S2 shows the top 5 most differentially weighted genes in each group, and we conducted reactome enrichment analysis on the five genes in the HCM disease (Fig. 6d and Supplementary Table S3), identifying five related pathways. Of these, pathways 1 and 3 are related to the Notch1 gene, further highlighting Notch1’s essential role in embryonic development and heart formation and its role as a crucial factor in the proliferation and differentiation of cardiac stem cells [31]. Inhibition of Notch1 signaling could lead to life-threatening dilated cardiomyopathy, emphasizing that scKEPLM can not only validate known disease-related genes and pathways but also expand our perspectives in researching cardiac disease biomarkers and therapeutic targets.

Fibroblasts play an essential role in the heart, responsible for synthesizing the extracellular matrix (ECM) and collagen, ensuring the normal structure and function of the heart [32]. In cardiac diseases such as HCM and DCM, abnormalities in fibroblast function are closely related to pathological myocardial fibrosis, which leads to damage to the structure and function of the heart. We analyzed the gene expression of fibroblasts and presented a weight analysis of these cell populations in Fig. 6e and Supplementary Table S4. The results showed that, in the fibroblasts of patients with HCM, the top ten upregulated genes are mainly concentrated in the regulation pathway of the CDC42 GTPase activity cycle. Interestingly, these findings have been verified in a recent study [33]. The research indicates that CDC42 is related to the pathogenesis of HCM, and experiments based on mouse models have shown that the specific knockout of the CDC42 gene in the heart under stress leads to more significant cardiac hypertrophy. Two weeks and eight weeks after stimulation, these mice progress to heart failure at a faster rate.

At the same time, previous research observed that in DCM and HCM hearts compared to NF hearts, the expression of markers for certain endothelial and lymphoid cells, such as KIT, NRP2, and KLRF1 [29], was significantly increased. By analyzing these two cell populations, we found that the average expression levels of the previously reported marker genes increased in the respective cell types Fig. 6f-g, consistent with earlier study outcomes, confirming that scKEPLM can effectively identify relationships related to genes in the context of disease. By delving deeper into scKEPLM’s predictions, we have captured not only the changes in marker genes but also unveiled potential mechanisms behind the progression of the disease. Particularly in cardiomyopathy model studies, genes and related biological pathways highlighted by the model’s recognized Gaussian weights not only reflect the close relationship between specific cardiac diseases and specific genes and biological processes but also emphasize the importance of heterogeneity at the cellular level and the complexity of intercellular communication in the development of heart diseases. Therefore, the scKEPLM model provides a powerful tool that can deepen our understanding of the mechanisms behind heart failure and other cardiac diseases and offer valuable insights for the future diagnosis and treatment of heart diseases.

#### 1.5.2 scKEPLM captures cell annotation markers

After fine-tuning the scKEPLM model on the Muraro dataset for cell type annotation[34], we utilized a Gaussian attention mechanism to distill the unique gene expression features of each cell type. Then we performed differential analysis through generated weight coefficients instead of directly comparing gene expression levels. It enables us to identify key genes related to specific cell types, rather than merely comparing their levels of expression.

It is noteworthy that the scKEPLM model successfully identified key genes in Alpha-cells, including well-recognized ones such as IRX2 and LOXL4 [35]. Such results further confirm previous findings about these markers, showing the ability of the scKEPLM model to identify potential cell marker genes.

Moreover, in the enrichment analysis of the top 30 genes of interest to the model in Alpha-cells, by comparing records across different gene databases, we found these genes showed higher enrichment levels in categories related to their corresponding cellular functions. For instance, as shown in Extended Data Fig. 5 and and Supplementary Table S5-6, the gene clusters specifically related to the functionalities of Alpha-cells were particularly prominent in the enrichment analysis. These analysis outcomes demonstrate the efficacy of the scKEPLM model in capturing fundamental cell characteristics during the training process.

## 2. Discussion

We developed scKEPLM, a knowledge-enhanced single-cell transcriptome pre-trained model. scKEPLM is the first single-cell foundation model to integrate genetic knowledge graph pre-training and RNA sequence pre-training into a unified framework. The knowledge graph enhancement enables scKEPLM to capture extensive interactions between independent genes and dynamic network information within cells, allowing it to simultaneously understand and learn both the biological semantics and genetic contexts of genes. scKEPLM innovatively employs the Gaussian attention mechanism within the transformer architecture to align key knowledge with RNA sequence representations, substantially enhancing its interpretability in learning gene and cell profiles.

scKEPLM demonstrates exceptional generalization capabilities across a variety of downstream tasks, including gene annotation, cell and cell subtype annotation, cancer drug response prediction, and biomarker identification. First, scKEPLM comprehensively outperforms existing state-of-the-art single-cell transcriptome models in all 12 downstream tasks. Secondly, a series of domain transfer experiments, including cross-tissue and cross-platform cell annotation and drug response prediction from single-cell to bulk data, prove the robust transferability of scKEPLM. In addition, scKEPLM recognizes the importance of genes in cell annotation by modeling gene expression in cells using a Gaussian attention mechanism. Through differential gene enrichment analysis of model weights, we discovered that scKEPLM is capable of distinguishing between diseased groups and identifying differential genes across various cell types. This capability emphasizes the key role of scKEPLM in understanding biological processes, clarifying disease mechanisms, and promoting innovative treatment strategies.

Ablation research shows that the performance of the single-cell transcriptome pre-training model is significantly enhanced by incorporating genetic knowledge. This enhancement not only effectively aggregates marker genes in single cells but also prioritizes them in disease-related research, thus enhancing the model’s data interpretation ability. During the pre-training phase, scKEPLM effectively tackles challenges related to the modeling of gene interactions and the optimization of masking strategies within the Transformer framework, signifying substantial advancements in the domain of single-cell data analysis.

In summary, the scKEPLM presented in this paper is an novel single-cell transcriptome pre-training model that leverages effective knowledge enhancement and self-supervised learning strategies. It exhibits exceptional performance and interpretability across a wide range of downstream tasks at the gene and cellular levels. Our previous experiments demonstrate that single-cell transcriptome pre-training models can be enhanced by incorporating a broader range of multi-modal information. In future research, we will try to integrate additional prior knowledge, including literature annotations, gene sequences, and epigenetic modifications, in order to develop a comprehensive single cell model and benefits from multimodal collaboration.

## 3. Methods

### 3.1 scKEPLM architecture and pretraining

The scKEPLM model consists of three primary components. The first involves knowledge construction and representation. This phase utilizes additional biological knowledge to construct a knowledge graph, followed by a knowledge embedding method to learn the knowledge representation. The second component is scKEPLM pretraining. This process draws an analogy between single-cell RNA-seq data and natural language. Gene expression sequences from individual cells are treated as sentences, with each gene or transcript akin to a word in this biological language. The expression levels of genes represent the “meaning” of these sentences. scKEPLM incorporates two parallel Masked Language Models (MLMs) [36]: the sequence encoder (S-Encoder) and the knowledge encoder (K-Encoder), aligning with the conventional BERT paradigm. Within the encoder module, each unit features a self-attention mechanism and a feed-forward neural network. The Gaussian cross attention mechanism integrates additional biological knowledge into the sequence information, effectively mapping the two representations into a unified feature space. It is designed to reduce feature redundancy, enhance weight differentiation among genes, and highlight crucial information in the input sequence, thereby reducing computational complexity [37]. It also prevents the oversmoothing of embeddings, particularly in cases of longer input sequences. The MLMs train the model to predict masked or missing labels in the sequence, focusing on understanding the contextual relationships between genes and inferring missing expressions.

#### 3.1.1 Knowledge construction and representation

Domain knowledge in biology is essential for enhancing large pre-trained models in single-cell RNA sequencing. We incorporate the STRING database (Search Tool for the Retrieval of Interacting Genes/Proteins) as an external knowledge graph. The STRING database offers a comprehensive protein-protein interaction (PPI) network, crucial for understanding cellular protein interactions. It allows pre-trained models to identify significant protein interactions in single-cell RNA sequencing analysis. Additionally, STRING enhances biological pathway analysis, enabling the identification of pathways and functional modules within gene expression data. These capabilities provide deeper insights into cellular states at the single-cell level, where distinct gene expression patterns are observed. The interaction data from STRING are also vital for identifying biomarkers and genes associated with specific diseases or biological traits, playing a key role in disease diagnosis and treatment planning.

To fully utilize the fine-grained semantic relationships between genes in the existing knowledge base, we use the STRING [6] (Search Tool for the Retrieval of Interacting Genes/Proteins) database to acquire external knowledge. STRING is a powerful database for studying protein-protein interactions. It provides data on established and predicted protein interactions, including both direct (physical) and indirect (functional) connections [6].

In the knowledge graph 𝒢 = (ℰ, *ℛ, 𝒯*), ℰ represents the set of gene entities, ℛ denotes the set of relations, and 𝒯 comprises triples (*h, r, t*). Each triple indicates that the head gene entity *h* is related to the tail gene entity *t* via relation *r*. For each gene entity *e*, its initial knowledge embedding is transformed from a one-hot vector to a *d*-dimensional vector **e** using a transformation matrix **W** *∈* ℝ^*V ×d*^, where *V* is the complete vocabulary containing all gene tokens. When *e* acts as the head entity *h*, **h** = **e**, and when *e* acts as the tail entity *t*, **t** = **e**. For each gene relation *r*, we introduce a transformation matrix **M**^(*r*)^ *∈* ℝ^*d×k*^. This matrix captures the specific characteristics of the relation and projects entity embeddings **h** and **t** from the entity space to a relation-specific space, as illustrated by the following equation:

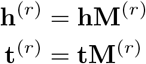

We then define a scoring function to evaluate the possibility of a triple (*h, r, t*) using the L1 or L2 distance:

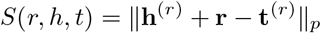

where the representation of head entity gene *h* and the tail entity gene *t* are both transformed into the relation space *r*, ℝ^*k*^.

To learn the knowledge embeddings of genes and transformation matrices, we use a margin-based loss function, to encourage the model to assign higher scores to true triples (*h, r, t*) and lower scores to corrupted triples (*h, r, t*^*I*^), where *t*^*I*^ represents a negative sample. Finally, we obtain the updated *d*-dimensional vector **e** as the initial knowledge representation for each gene, which can then be sequentially input into the knowledge encoder.

#### 3.2.1 Sequence encoder and knowledge encoder

scKEPLM includes a sequence encoder and a knowledge encoder, both adhering to the masked language model paradigm. Specifically, for a given gene sequence from a single cell *{g*_1_, …, *g*_*n*_*}* and its corresponding entity sequence *{e*_1_, …, *e*_*n*_*}*, where *n* is the length of the input RNA-sequence, the sequence encoder first combines the token embedding and positional embedding for each gene to create its input embedding. It then computes the syntactic features *{***s**_1_, …, **s**_*n*_*}* as follows:

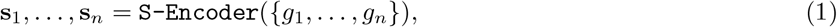

Similarly, we take the representation of the knowledge entity *{e*_1_, …, *e*_*n*_*}* learned from the pre-trained TransR model as the input, and through the knowledge encoder K-Encoder, we obtain the knowledge representation of the gene.

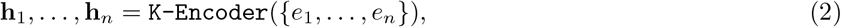

where both S-Encoder(·) and K-Encoder(·) are multi-layer bidirectional Transformer encoders.

After processing the gene sequence through the two encoders, we obtain the contextual representation matrix 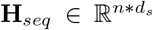 and knowledge representation matrix 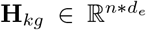, where *d*_*s*_ and *d*_*e*_ are hidden dimension of S-Encoder and K-Encoder.

#### 3.1.3 Gaussian cross attention mechanism

To fuse sequence and knowledge representations more effectively, we propose a Gaussian cross-attention mechanism. Traditional cross-attention relies on Q, K, and V mappings with softmax weight normalization, which can result in overly smoothed weights or challenges in capturing long-range correlations, especially with long sequences. To address this, we use Gaussian normalization in place of softmax normalization to better capture the strong correlation between knowledgeable and contextual information [37, 38]. The details of Gaussian cross-attention are as follows:

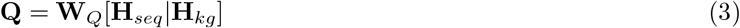

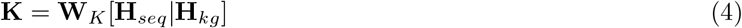

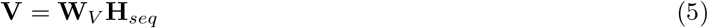

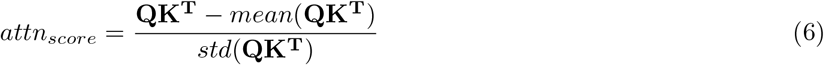

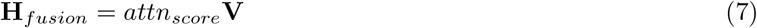

where the weight matrices **W**_*Q*_, **W**_*K*_ and **W**_*V*_ are learnable matrices, and [*·*|*·*] represents the vector concatenation operation. The functions *mean*(·) and *std*(·) are employed for calculating the mean and standard deviation. **H**_*fusion*_ denotes the embedding matrix following Gaussian cross-fusion.

#### 3.1.4 Pre-training and fine tuning

Overall, scKEPLM is divided into two stages: pre-training and fine-tuning for downstream tasks. During the self supervised pre-training, a certain percentage (usually 15%) of tokens in each sequence is randomly selected and replaced with a [mask] token. Then, the modified sequence and its masked tokens are fed into a multi-layer transformer encoder. Each Transformer layer captures contextual information from both directions (left and right context) around each token. It then predicts the original values of the masked tokens based on their context. While this prediction is not used directly in sequence encoding but helps the model learn better representations during pre-training. The objective function for the model’s pre-training process is as follows:

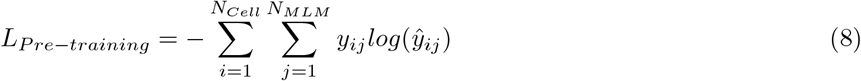

where *N*_*Cell*_ represents the total number of cells, *N*_*MLM*_ denotes the number of masked genes in the masked language model, and *y*_*ij*_ and *ŷ*_*ij*_ represent the ground-truth and prediction results for the genes, respectively. The optimal hyperparameters during pre-training include an initial learning rate of 10^−3^, using the AdamW optimizer, with the weight decay set at 10^−10^.

### 3.2 Dataset

In our study, we introduce an advanced models scKEPLM that excels in analyzing single-cell transcriptome data through two key phases: pre-training and fine-tuning. In the pre-training phase, we utilize a vast corpus of unlabeled data for unsupervised learning to develop deep representations. We meticulously compiled over 41 million human single-cell transcriptome data points from 569 publicly available datasets, conducting statistical analyses to select 25,424 genes with the highest representativeness for further analysis. Concurrently, we incorporate six types of relationships from the STRING [6] database to construct a gene interaction graph. The knowledge graph furnishes our model with extensive prior information, thereby enhancing its comprehension of gene interactions. Notably, our knowledge graph covers approximately 75% of genes, ensuring the model’s thorough and precise understanding of gene interactions during the fine-tuning phase. In this phase, we specifically fine-tune the model using different datasets tailored to individual tasks, further enhancing its performance.

#### 3.2.1 The Panglao dataset - Embryonic Stem Cells

The dataset was downloaded from the PanglaoDB [13] website (https://panglaodb.se/) and comprises humancultured Embryonic Stem Cells (ECs) data with the accession number SRA553822-SRS2119548, encompassing a total of 34,254 cells. In our study, we meticulously selected 15,000 cell samples from this dataset sourced from PanglaoDB’s Embryonic Stem Cells data. This randomly chosen subset of samples was employed for predicting genes associated with bivalent unmethylation and exclusive lysine methylation.

#### 3.2.2 Dataset for iPSC to Cardiomyocyte Differentiation

The dataset [15] originates from a iPSC to cardiomyocyte differentiation study available in the Gene Expression Omnibus (GEO) database. The data were generated using parallel profiling on Drop-seq or DroNc-seq methods. This publicly accessible dataset encompasses four distinct states: iPSC, Cardiac Progenitor, Cardiomyocyte 1, and Cardiomyocyte 2, totaling 70,060 cells. Our research focuses on two primary tasks: first, a long and shortterm gene classification task aimed at identifying and categorizing genes crucial in the iPSC to cardiomyocyte differentiation process. Second, a cross-platform cell annotation task, striving to provide more accurate annotations and classifications for each cell state by integrating data from multiple parallel profiling sources. The dataset can be downloaded from the GEO database under accession number GSE1290963.

#### 3.2.3 Muraro Dataset

Utilizing cel-seq2 sequencing technology, this dataset successfully extracted data from human pancreatic cells. The dataset [34] comprises a total of 1601 cells, covering four distinct cell types: alpha cells (843 cells), beta cells (445 cells), delta cells (203 cells), and gamma (PP) cells (110 cells). This dataset furnishes detailed information regarding gene expression levels in human pancreatic cells, offering a rich resource for further investigations into pancreatic function and cell type differentiation. The dataset is available for download on the GEO database under accession number GSE85241.

#### 3.2.4 Braon Dataset

The Braon dataset [39] successfully extracted human pancreatic cell data using the in Drop sequencing technology. This dataset comprises a total of 2,650 cells, encompassing four distinct cell types: alpha cells (912 cells), beta cells (1,243 cells), delta cells (339 cells), and gamma (PP) cells (156 cells). The dataset is available for download on the GEO database under accession number GSE84133.

#### 3.2.5 Segerstople Dataset

The Segerstople dataset [40] utilized SMARseq sequencing technology to extract data from human pancreatic cells. This dataset encompasses 1,656 cells, including four main cell types: alpha cells (1,008 cells), beta cells (308 cells), delta cells (213 cells), and gamma (PP) cells (127 cells). The dataset is available for download on the ArrayExpress database under accession number E-MATB-5061.

#### 3.2.6 Zheng68k Dataset

The dataset [41] is a widely used public resource in single-cell RNA sequencing research, primarily employed to investigate gene expression in human peripheral blood mononuclear cells (PBMCs). It comprises a total of 68,482 cells, covering 13 different cell subtypes, specifically: CD8+ Cytotoxic T (20,773 cells), CD8+/CD45RA+ Naive Cytotoxic (16,666 cells), CD56+ NK (8,776 cells), CD4+/CD25 T Reg (6,187 cells), CD19+ B (5,908 cells), CD4+/CD45RO+ Memory (3,061 cells), CD14+ Monocyte (2,862 cells), Dendritic (2,099 cells), CD4+/CD45RA+/CD25-Naive T (1,873 cells), and CD34+ (277 cells). Detailed information about this dataset can be downloaded from the 10X Genomics website (https://www.10xgenomics.com/resources/datasets).

#### 3.2.7 Heart Disease Dataset

The dataset [29] consolidates cellular data from 42 patients, including 12 with Dilated Cardiomyopathy (DCM), 15 with Hypertrophic Cardiomyopathy (HCM), and 16 with Non-failing hearts (NF), amounting to a total of 579,159 cells. To acquire single-cell RNA sequencing data, we employed the 10X Genomics technology. Specifically, the cell counts for patients with Hypertrophic Cardiomyopathy, Non-failing hearts, and Dilated Cardiomyopathy were 230,652, 182,317, and 166,190, respectively. Detailed information about the data can be found at https://singlecell.broadinstitute.org/single_cell/study/SCP1303.

#### 3.2.8 Cancer Drug Response Dataset

The cancer drug response dataset [28] integrates two databases by retaining shared genes, as well as all cell lines and drugs. In total, the dataset has collected extensive drug response data for 1,280 cancer cell lines and 1,557 drugs/compounds, as well as their expression profiles across 15,962 genes. Detailed data can be found at https://github.com/OSU-BMBL/scDEAL.

### 3.3 Main comparison methods

#### 3.3.1 Geneformer

Geneformer [42] is a large-scale context-aware pretrained language model designed for transcriptomic data. It aims to make predictions in network biology with limited data through transfer learning. Geneformer utilizes recent advancements in self-attention to maintain focus on the extensive input space of genes expressed in each single-cell transcriptome. It understands which genes are most crucial, optimizing prediction accuracy for a given learning objective. In this paper, it is used as a baseline model.

#### 3.3.2 scGPT

scGPT [43] is a foundation model for single-cell biology based on a generative pre-trained transformer that utilizes a database of over 33 million cells. To handle the large-scale data, scGPT employs in-memory data structures to store hundreds of datasets for rapid access and has established a unified generative pre-training workflow, specifically tailored for non-sequential genomic data to learn cell and gene representations simultaneously. In this paper, it is used as a baseline model.

#### 3.3.3 scFoundation

scFoundation [10] is a large-scale pre-trained model that utilizes the xTrimoGene architecture with 100 million trainable parameters. This model is trained on human single-cell transcriptomic data, which includes highthroughput observations of complex molecular features in all known types of cells. scFoundation is considered a large-scale model in terms of the size of its trainable parameters, the dimensionality of genes, and the number of cells used for pre-training. In this paper, it is used as a baseline model.

#### 3.3.4 scBERT

scBERT [21] is a single-cell embedding model based on BERT (Bidirectional Encoder Representations from Transformers), designed for learning cellular representations within single-cell RNA sequencing data. Leveraging deep learning techniques, particularly the pre-trained BERT model, it aims to capture intricate relationships and features within gene expression data. In this paper, it is used as a baseline model.

#### 3.3.5 CasTLe

CasTLe [44] (Cell-type Annotation by Transfer Learning) is a single-cell annotation method that employs the concept of transfer learning. This approach guides the prediction of labels for unknown cells in a target dataset by leveraging information learned from a known dataset (source domain). The unique aspect of CasTLe lies in its ability to transfer cell types across different datasets, enabling the annotation of unlabeled cells in the target dataset. This capability proves particularly useful for handling single-cell data from diverse sources or conditions.

#### 3.3.6 scDeepsort

scDeepsort [20] is a single-cell annotation method that integrates deep learning and supervised learning. It is based on deep neural networks, allowing it to annotate cell types in new single-cell data by learning features and cell type labels from extensive single-cell RNA sequencing data. Leveraging deep learning techniques, scDeepsort excels in capturing complex cell type features during the learning process, contributing to its outstanding performance in cell annotation tasks. Additionally, it demonstrates adaptability to large-scale datasets, making it valuable for handling diverse and abundant single-cell RNA sequencing data.

### 3.4 System Analysis

#### 3.4.1 Gene Annotation

To evaluate scKEPLM’s effectiveness in gene annotation, we fine-tuned the pre-trained scKEPLM model and compared it with three machine learning based methods and one advanced pre-training model. Initially, we ranked and sorted the target single-cell expression data based on rank-value criteria, selecting the top 2048 genes to represent cell features effectively. These genes were then converted into gene sequences, serving as input for the model. For gene annotation, scKEPLM was fine-tuned, specifically by adding one or two MLP layers for classification tasks. It’s important to note that we froze the layers of scKEPLM, demonstrating that excellent gene annotation performance can be achieved by only fine-tuning the one or two MLP layers added subsequently.

#### 3.4.2 Cell Annotation

To evaluate the scKEPLM model’s proficiency in cell annotation, we compared it against twelve conventional methods. The process involves inputting single-cell gene tokens into the pre-trained scKEPLM model and using the average of all gene embeddings within a single cell as the cell’s representative embedding. During the finetuning phase, we enhanced scKEPLM with a multilayer perceptron (MLP), adjusting both the MLP and the last two layers of the scKEPLM unit. The adjustment is crucial, as cell-level annotation tasks often deal with significant heterogeneity and complex behaviors, requiring a deeper network structure to discern differences and relationships between cells. Moreover, to test the model’s adaptability across different domains, we fine-tuned it using data from the Drop-seq platform and evaluated it on sequences from the DroNc-seq. Our findings affirm the model’s consistent effectiveness across diverse tasks.

#### 3.4.3 Model Interpretability

By constructing a Gaussian weight matrix, we generate a matrix representing the mutual importance of genes for each sequence. Initially, we average the weight matrix of the three-layer scKEPLM model to obtain a comprehensive weight representation. Subsequently, we further average along the columns to derive a vector representing the relative importance of each gene in the cell. We utilize this vector as a substitute for gene expression to explore the interpretive approach of the model, including gene difference analysis and functional enrichment analysis. Our model demonstrates remarkable ability, not only in identifying genes with significant differences in expression but also in revealing key biological activities. Importantly, when comparing patients with healthy individuals, this model successfully identifies genes as disease markers and accurately highlights the characteristics of their abnormal expression. Furthermore, we explore capturing gene associations without fine-tuning the model. The research indicates that the model effectively captures interactions between landmark genes. Through these findings, our model not only provides high-quality interpretation of biological information but also offers valuable tools and insights for biomedical research.

## Acknowledgments

We thank the anonymous reviewers for their constructive suggestions. This work was partially supported by the National Science Fund for Distinguished Young Scholars [62225109] and the National Natural Science Foundation of China [62276059, 62072095]. Corresponding author: Guohua Wang.

## Data Availability

All data utilized in this study is publicly available, and its usage is thoroughly explained in the methods section. The Panglao dataset was obtained from https://panglaodb.se/. The published Zheng68K dataset was downloaded from https://support.10xgenomics.com/single-cell-gene-expression/datasets (SRP073767) under the “Fresh 68K pbmc” section. The nine datasets for single-cell annotation across different human tissues can be downloaded from Hugging Face at https://huggingface.co/datasets/ctheodoris/Genecorpus-30M. The pancreas dataset was obtained from the GEO database at https://www.ncbi.nlm.nih.gov/geo/ (Baron: GSE84133, Muraro: GSE85241, Xin: GSE81608, MacParland: GSE115469). The Segerstolpe dataset was downloaded from the ArrayExpress database at https://www.ebi.ac.uk/arrayexpress (ID: E-MTAB-5061). Marker genes for cells were obtained from https://xteam.xbio.top/CellMarker/. Additionally, this paper provides all processed data in https://huggingface.co/datasets/catly/scKEPLM-41M.

## Code Availability

All relevant data and source codes including the the pre-trained model, can be downloaded from https://huggingface.co/catly/scKEPLM.

## Author contributions

G.W. conceived the work, assisted in algorithm design and implementation. Y.L. and G.Q. contributed to algorithm design, implementation, and computational experiment analysis. Y.L. drafted the initial manuscript. G.W. and Y.L. contributed to manuscript revision. G.Q. contributed to data acquisition and processing.

## Competing interests

All submitting authors declare no competing interests.

---

**Extended Data Fig. 1:**
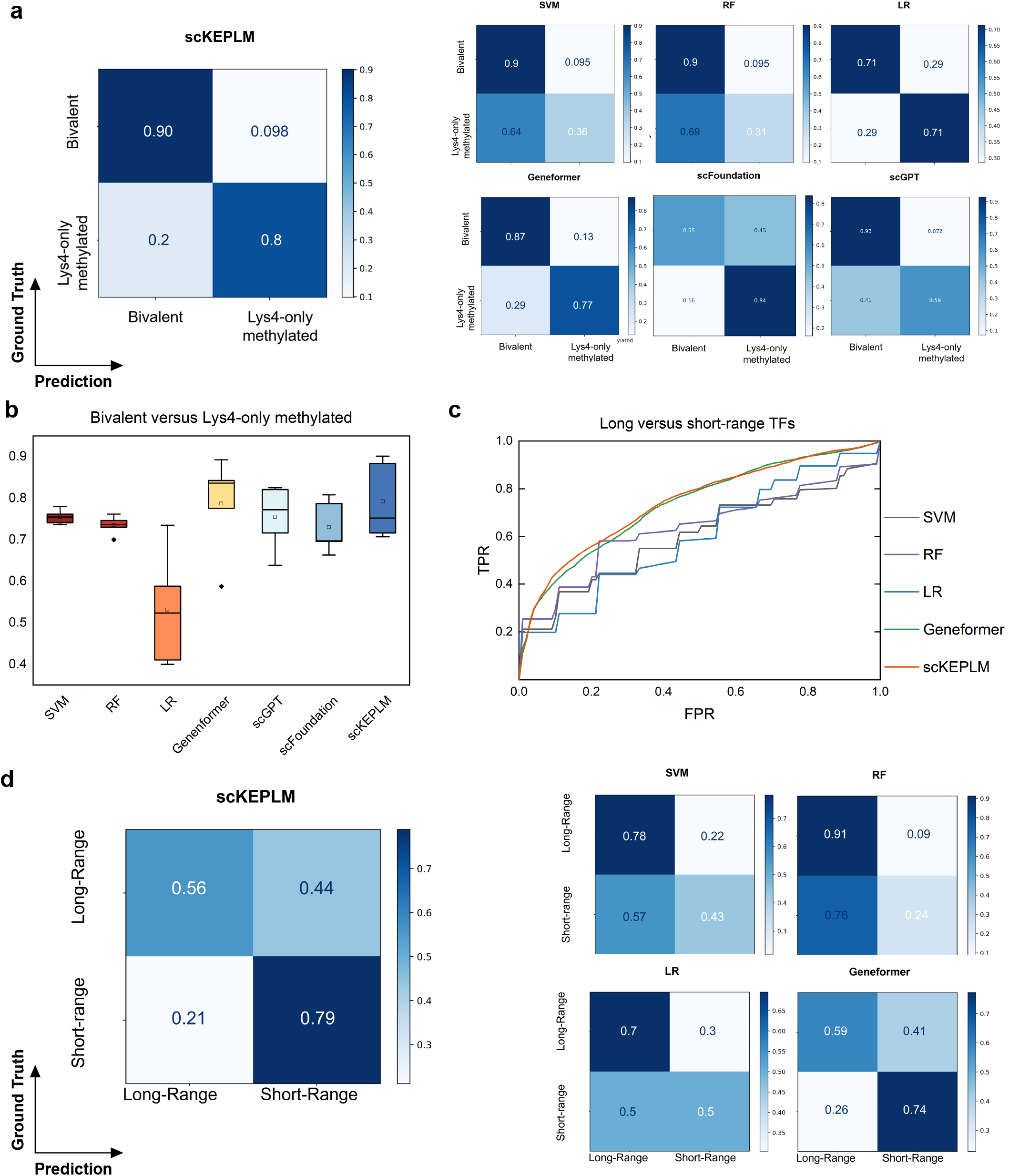
scKEPLM for efficient gene annotation. **a**, Confusion matrices of the six comparision methods and scKEPLM model for bivalent and Lys4-only Methylation prediction. **b**, Comparison of bivalent and Lys4-only Methylation prediction F1-score using box plots. **c**, Comparison of ROC curves for long and short transcription factor prediction. **d**, Confusion matrices of the four comparision methods and scKEPLM model for long and short transcription factor prediction.

**Extended Data Fig. 2:**
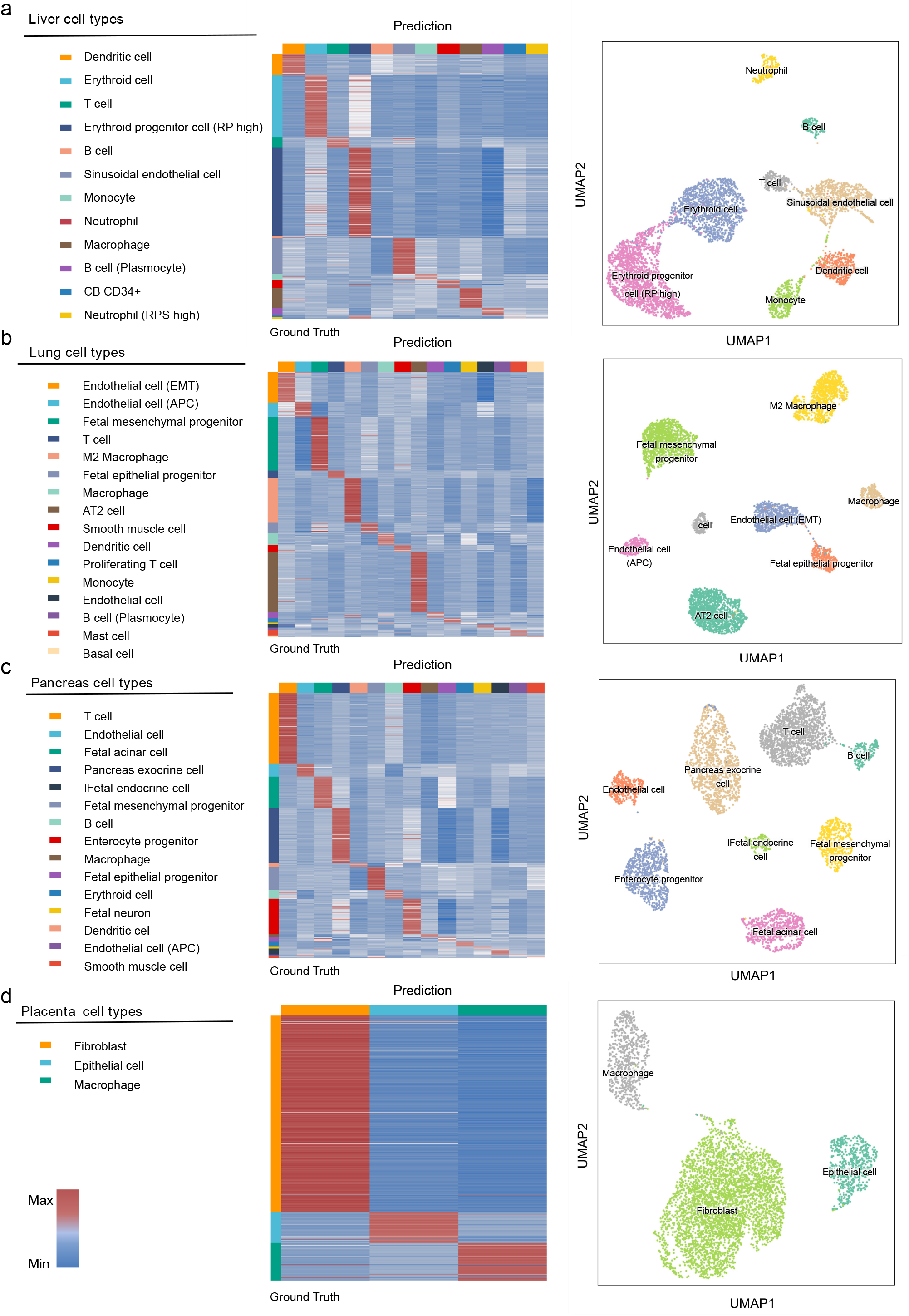
scKEPLM for efficient cell annotation with limited data. **a**, Liver, **b**, Lung, **c**, Pancreas, **d**, Placenta out of sample predictions by scKEPLM fine-tuned to distinguish cell types in each tissue (training on 80 % of cells, predictions on held-out 20% of cells).

**Extended Data Fig. 3:**
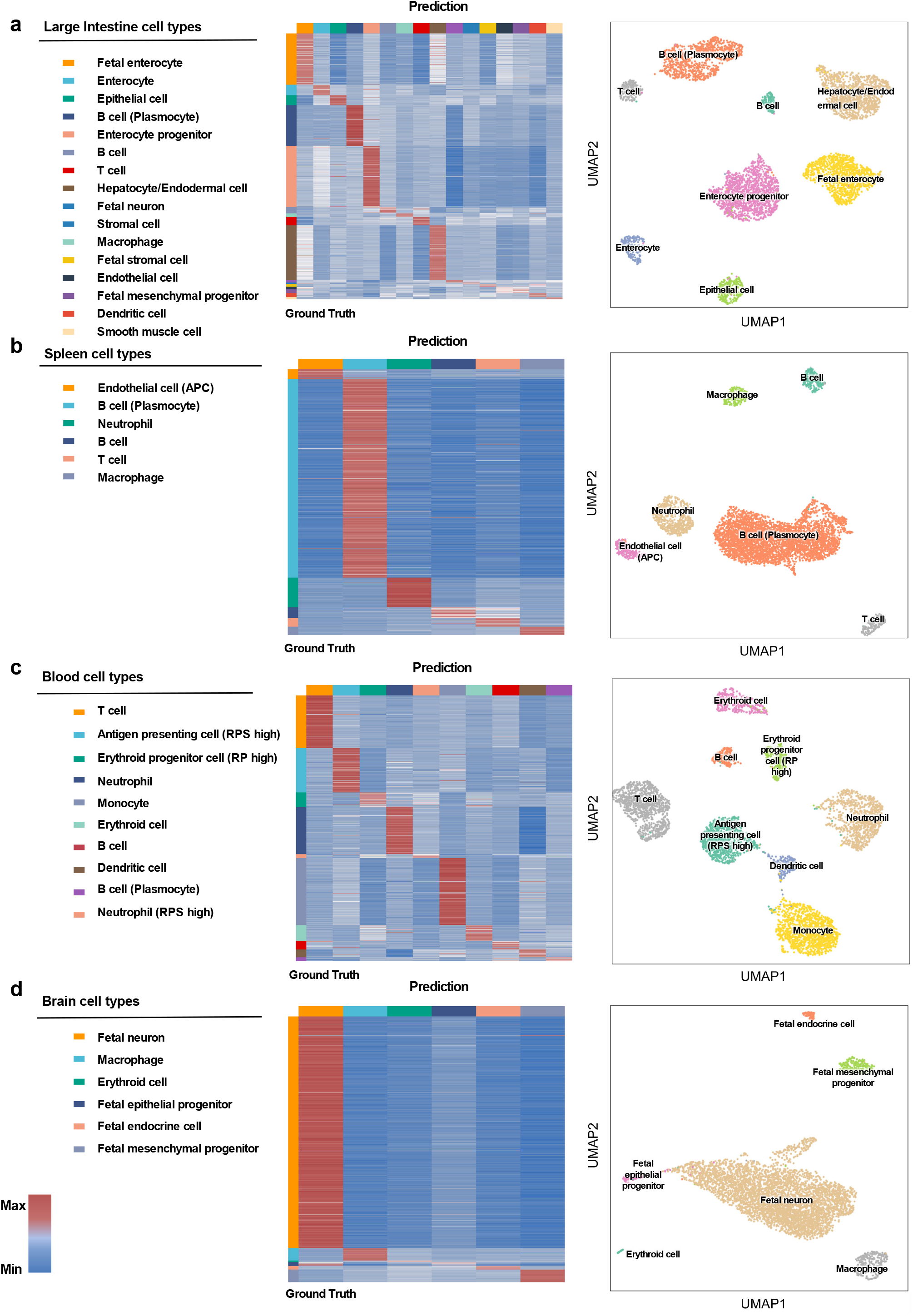
scKEPLM for efficient cell annotation with limited data. **a**, Large Intestine, **b**, Spleen, **c**, Blood, **d**, Brain out of sample predictions by scKEPLM fine-tuned to distinguish cell types in each tissue (training on 80 % of cells, predictions on held-out 20% of cells).

**Extended Data Fig. 4:**
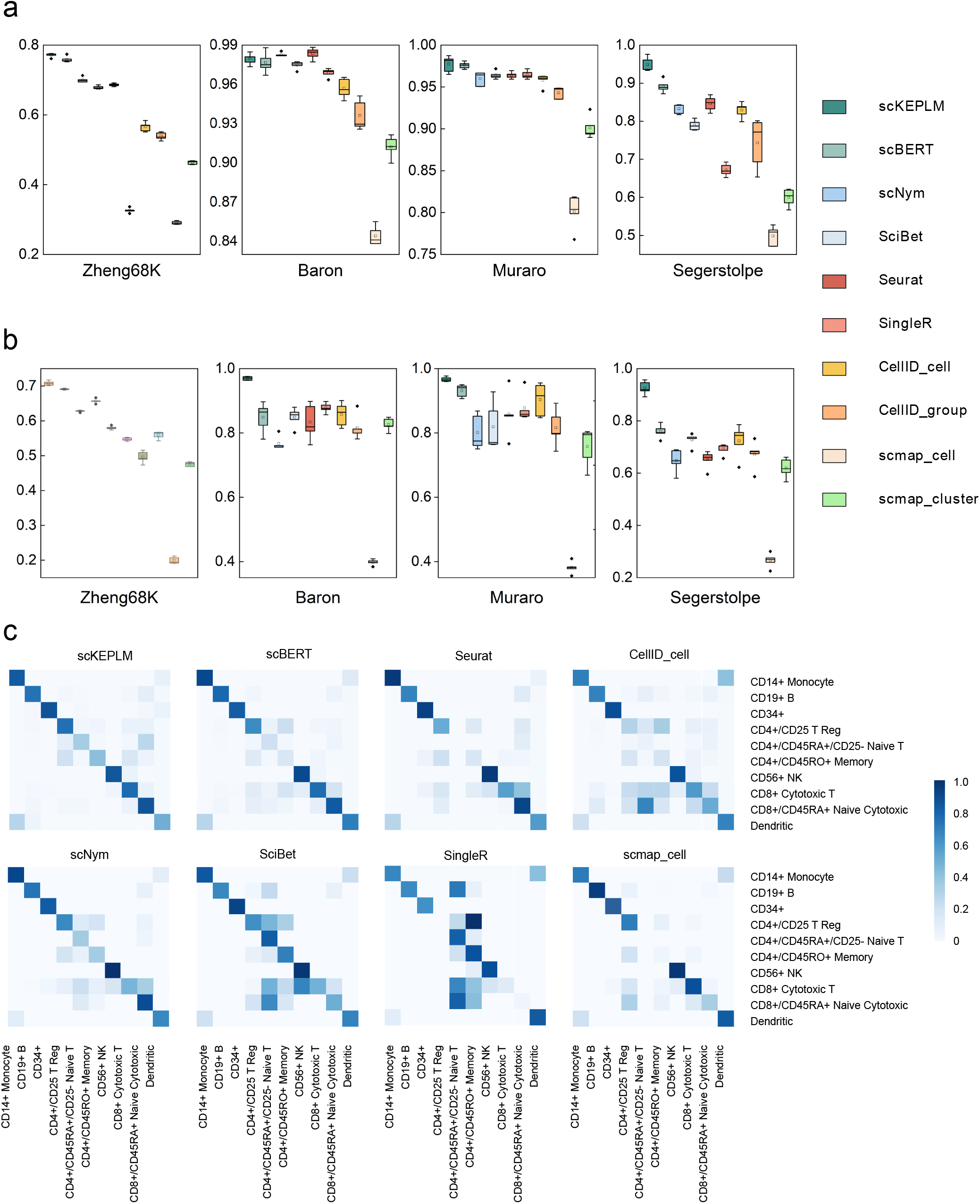
Performance of scKEPLM and other methods under cell annotation. **a**, The boxplots compare the performance of different models across various datasets. The boxplots display the distribution of ACC for each model on a given dataset. **b**, The boxplots display the distribution of F1-score for each model on a given dataset. **c**, Confusion matrix heatmaps for cross-validation results of others baselines on the Zheng68K dataset. The confusion matrices for other methods are included in the supplementary data.

**Extended Data Fig. 5:**
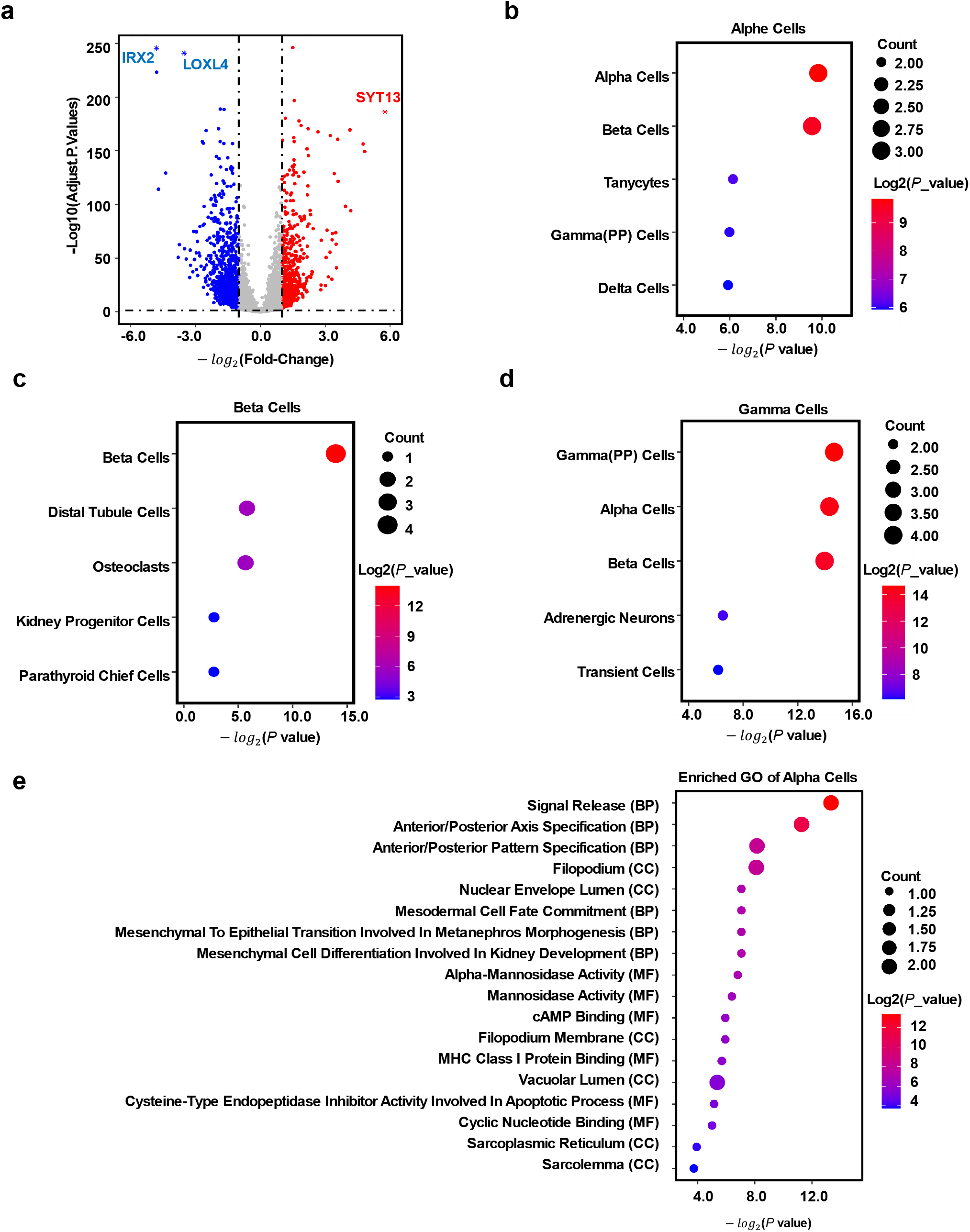
scKEPLM captures cell annotation markers. **a**, The weight obtained from alpha cells through scKEPLM is used as the expression value for differential analysis in the volcanic map, where the labeled genes are the marker gene of alpha cell. **b**, The enrichment analysis results of the top ten genes in alpha cells. **c**, The enrichment analysis results of the top ten genes in beta cells. **d**, The enrichment analysis results of the top ten genes in gamma cells. **e**, Enrichment pathway analysis results of the top thirty most concerned genes in alpha cells.

## Notes

https://huggingface.co/catly/scKEPLM

### Competing Interest Statement

The authors have declared no competing interest.

